# Using experimental data as a voucher for study pre-registration

**DOI:** 10.1101/213439

**Authors:** Matan Mazor, Noam Mazor, Roy Mukamel

## Abstract

Undisclosed exploitation of flexibility in data acquisition and analysis blurs the important distinction between exploratory and hypothesis-driven findings and inflates false-positive rates^1–4^. Indeed, recent replication attempts have revealed low levels of replicability, pointing to high rates of false-positives in the literature^5–10^. A contemporary solution to this problem is pre-registration: commitment to aspects of methods and analysis before data acquisition^11^. This solution is valid only to the extent that the commitment stage is time-locked to precede data collection. To date, time-locking can only be guaranteed by introducing a third party such as peer reviewers at an early stage, making this solution less appealing for many^12^. Here we adapt a cryptographic method^13^ to encode information of study protocol within random aspects of the data acquisition process. This way, the structure of variability in the data time-locks the commitment stage with respect to data acquisition. Being independent of any third party, this method fully preserves scientific autonomy and confidentiality. We provide code for easy implementation and a detailed example from the field of functional Magnetic Resonance Imaging (fMRI).

---

Pre-registration of study plans prior to data collection sharpens the distinction between hypothesis-driven and exploratory phases of the scientific process^11,14^. The importance of this distinction has been recently highlighted following numerous replication failures in the life and social sciences ^5–10^. Nonetheless, many researchers avoid pre-registration out of concern for their scientific autonomy and confidentiality^12^: involving an external party at an early stage of work might introduce delays to study commencement, make researchers more dependent on the publishing agency, or expose them to the risk of being scooped.

Here we introduce a pre-registration scheme that is inspired by cryptographic protocols^13^ and is performed in-lab, without the involvement of any third party. This scheme guarantees that the study protocol has been specified before data acquisition, making the registration time-locked — an essential feature for its validity. In order to fully maintain scientific confidentiality and autonomy, we break the pre-registration process to a commitment stage, performed by the researcher prior to data acquisition, and a verification stage that can be performed by anyone at any later stage. This division alleviates the need for an external inspector to time-lock the registration, and relegates the vouching process to the structure of variability of the data.

The scheme exploits random features in the experimental design to time-lock study plans. It is therefore applicable for experiments with aspects that can be determined in a pseudorandom fashion (such as the timing, order, or type of experimental events). It is assumed that different randomizations would yield different patterns in the data, and that post-hoc manipulation of the data with the purpose of introducing an alternative randomization is detectable. These conditions are met in many experimental designs used across scientific fields including neuroscience and psychology, and for different data types, such as images, behavioral measures, cell recordings and functional neuroimaging data.

## commitment stage

1. Before data acquisition, a protocol file is saved to a *protocol folder* together with any available details to which the authors wish to commit (such as number of measurements, predictions and analysis parameters that will be used). A script that uses a pseudo-random number generator (PRNG) to determine the experimental random aspects is also saved to the same folder.
2. A cryptographic hash function is applied to the protocol folder. This results in a sequence of bits that for all intents and purposes is unique to the protocol folder (*protocol sum*).
3. The protocol sum is used as an initialization seed for the PRNG.
4. The PRNG is used to determine various random aspects of the experimental protocol, such as order and timing of events.
5. Upon publication, the protocol folder is uploaded to an online repository, and a link to this repository is included in the final manuscript. Raw experimental data is shared publicly, or made available upon request.

In practice, steps 2-3 can be performed by calling our preRNG function (Python, R, and Mat-lab implementations accompany the current manuscript) prior to using the pseudorandom number generator to determine the random aspects of the experiment. The preRNG function receives the protocol folder as argument (e.g., `preRNG`(’D:/experiment/protocolFolder.zip’). In cases where multiple randomization schemes are desired, the function can be called with an additional argument specifying the randomization serial number (see Appendix). In our implementations we used the SHA-256 hash function that outputs 256 bits for any arbitrary length input (NIST, 2002).

## verification stage

The commitment stage introduced a causal link between the acquired data and the content of protocol folder via a chain of dependencies.

1. The dependency of the acquired data on random components of the experimental design (red arrow 1 in figure 1) is a prerequisite for the use of this scheme.
2. The dependency of random components of the experimental design on the protocol sum (red arrow 2 in figure 1) was obtained through the initialization of the PRNG with the protocol sum as seed. This dependency is tight, since the behavior of the PRNG is deterministically set given a particular seed, and different seeds result in different behaviors^15^.
3. The dependency of the protocol sum on the protocol folder (red arrow 3 in figure 1) was obtained through the use of a cryptographic hash function. Such functions map arbitrary length inputs to sequences of bits of a fixed length such that finding a collision (i.e., two inputs that are mapped to the same sequence of bits) is infeasible.

Altogether, this chain of dependencies enforces a causal link and thus time-locks the acquired data with respect to the protocol folder. Due to this causal link, raw experimental data should be in line with the randomness incurred by the PRNG that has been initiated with the protocol-sum as seed. This can be verified by analyzing the shared data according to the analysis plans specified in the protocol folder, by visual inspection, or using any other data-based verification tests chosen by the verifier — be it an editor, a reviewer, or an interested reader.

**Figure 1:**
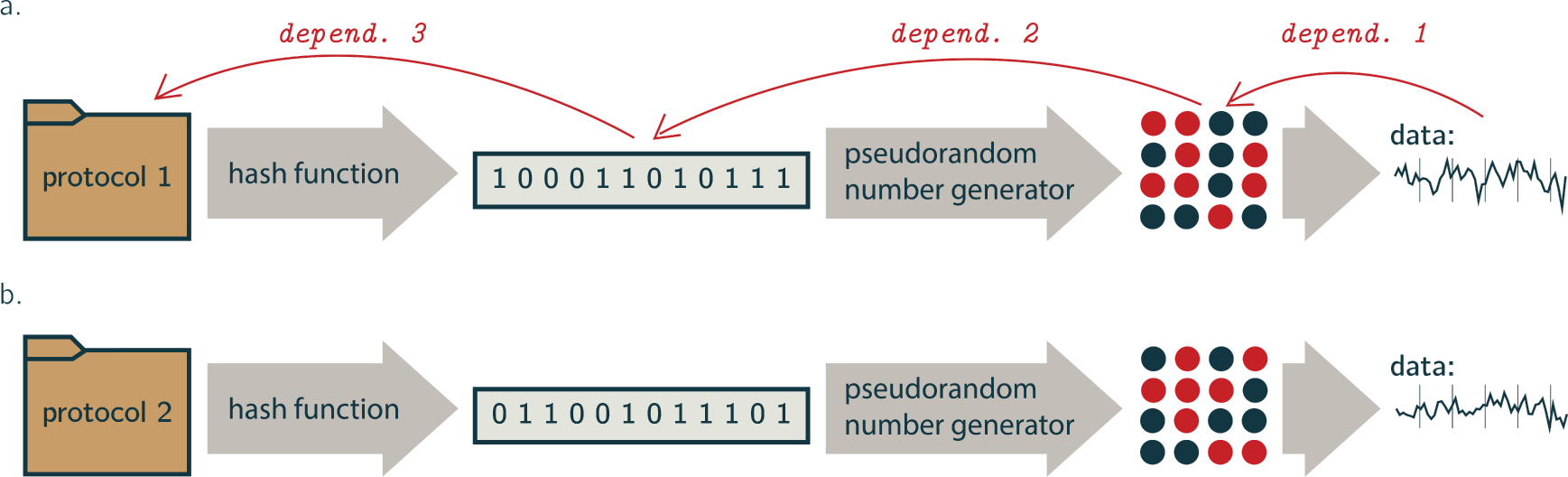
a. The pre-RNG scheme. Registration time-locking is obtained by making the acquired data (represented as line plots) dependent on the protocol folder via specific random components of the experimental design (represented as blue and red events). b. An alternative protocol folder results in a different randomization and therefore a different structure of data variability. This chain of dependencies time-locks the pre-registration with respect to data acquisition.

To demonstrate the use of the preRNG scheme we describe a hypothetical scenario involving a researcher (Alice) and an interested reader (Bob), based on an experiment that was conducted in our lab for the purpose of demonstration^16^. Alice examined cerebellar involvement in hand movement and committed to her study plans using preRNG. Bob wants to verify that certain findings that are especially relevant to his own research are hypothesis-driven, as reported. Alice’s paper and Bob’s verification are both included in the appendix to this paper.

Bob downloads the study protocol folder from the link provided in the manuscript and in it he finds a methods section specifying Alice’s choice to restrict her analysis to the cerebellum. Bob runs the preRNG function on the protocol folder, resulting in a protocol-sum that is identical to the one obtained by Alice (dependency number 3). Bob then uses the Python script that he found in the protocol folder to generate a pseudorandom sequence of experimental events, based on the resulting protocol-sum. Since Bob and Alice obtained an identical protocol-sum and since PRNGs are deterministic, Bob obtains the same sequence of events that was used by Alice in the actual experiment (dependency number 2). Given the high number of possible event orders in Alice’s experiment^16^, the likelihood of obtaining a particular sequence of events by chance is very small (< 10^−20^). Therefore the probability that a different PRNG seed would have resulted in a similar order of events is negligible.

In order to verify that the data reflects this randomization (dependency number 1), Bob writes to Alice and kindly asks for the raw experimental data. He then decides to perform a whole-brain GLM contrast between right and left hand movements, using the information he now acquired about the temporal order of events. Note that Bob is free to choose whatever verification analysis he finds fit and is not limited to the analysis reported by Alice (for additional verification steps he can use see Appendix). The resulting map is in line with Bob’s prior knowledge of robust lateralized brain activations in primary cortical areas. Since fMRI data are highly affected by the specific order of events, as reflected in the spatial and temporal dynamics of the signal, the alignment of the acquired data with the pseudorandom order of events is a reliable voucher for the pre-registration validity. Bob is now convinced that the randomization induced by the protocol-sum is in line with the data, and therefore that Alice’s specification of analysis plans in the protocol folder has been genuinely made prior to data acquisition.

The preRNG scheme allowed Alice to provide empirical support for her claim that certain choices have been made prior to data collection, without sharing her study plans with any external party at an early stage. This would not have been possible in any other pre-registration implementation. Unreviewed pre-registration^17^ platforms (UPR; aspredicted.org, osf.io) cannot guarantee that the registration of study protocol is indeed time-locked to precede data acquisition, as they only serve as an open vault for researchers to submit their study plans. To date, time locking of protocol registration can only be obtained by introducing an additional peer-review step at an early stage of work (reviewed pre-registration; RPR^18^). Knowing that reviewers might request changes in the experimental design reduces the incentive to pre-register studies for which data has already been collected. Some RPR schemes have the advantage of facilitating the publication of null results by committing to publish regardless of outcome — a feature that is not supported by our in-lab approach. Nonetheless, RPR compromises scientific confidentiality and autonomy, making this solution less appealing for many.

By providing a time-locked pre-registration scheme that maintains the confidentiality and autonomy of the scientific process, we hope to encourage research labs to pre-register predetermined aspects of their studies and by doing so delineate a clearer border between hypothesis-driven and exploratory findings. This is an important step in mitigating the contribution of undisclosed flexibility in data acquisition and analysis to the replicability crisis.

**Figure 2:**
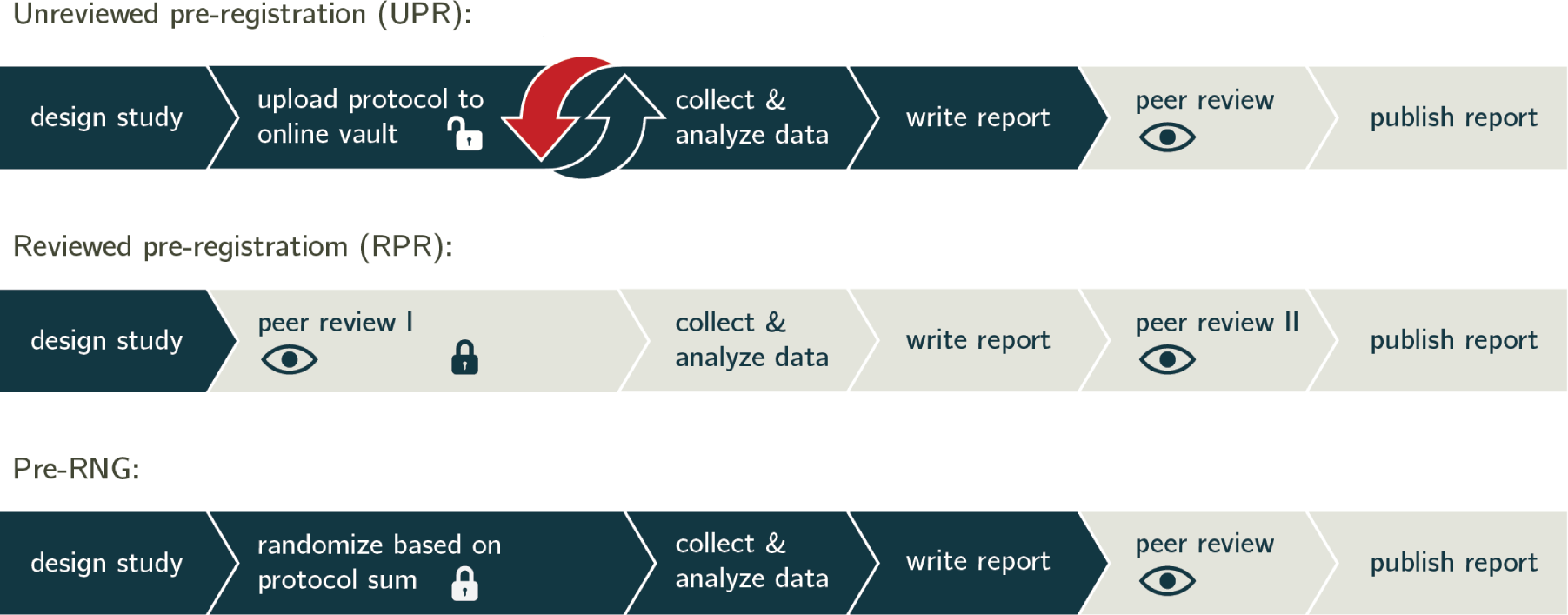
The three pre-registration schemes. The transition from dark blue to light gray indicates the first mandatory exposure of the research protocol to a third party. Lock icons represent the commitment to a specific research protocol. The red arrow in the UPR scheme represents the loophole allowing one to “pre-register” research plans even after data collection and exploration. Our pre-RNG scheme is time-locked but does not require early exposure.

## Acknowledgements

We thank Yoav Benjamini, Tal Galili, Tal Golan, Roni Maimon, Rafi Malach, Bats-Sheva Mazor, Jonathan Rosenblatt and Tom Schoenberg for their helpful insights and comments.

